# Dynamic stem cell states: naïve to primed pluripotency in rodents and humans

**DOI:** 10.1101/030676

**Authors:** Leehee Weinberger, Muneef Ayyash, Noa Novershtern, Jacob H. Hanna

## Abstract

The molecular mechanisms and signalling pathways that regulate the *in vitro* preservation of distinct pluripotent stem cell configurations, and their induction in somatic cells via direct reprogramming approaches, continue to constitute a highly exciting area of research. In this review, we provide an integrative synthesis on recent discoveries related to isolating unique naïve and primed pluripotent stem cell states with altered functional and molecular characteristics, and from different species. We overview pathways underlying pluripotent state transitions and interconversion *in vitro* and *in vivo.* We conclude by highlighting unresolved key questions, future directions and potential novel applications of such dynamic pluripotent cell states.

## Introduction

Pluripotency describes cells that have the potential to give rise to cells from all three embryonic germ-layers and possibly to primordial germ cells (PGCs), but not extra-embryonic tissues^1^. While pluripotency is a transient state *in vivo,* pluripotent cells can be derived from different stages of early embryonic development and indefinitely maintained in an artificially induced self-renewal state *in vitro,* by supplementing exogenous cues^2^. Thus, it is important to stress that self-renewal is not a defining feature of pluripotency and is only transiently assembled during early development. Pluripotency is highly dynamic and evolves at different stages of pre- and post-implantation stages^3^. However, the self-renewal aspect is a highly useful *in vitro* artificial “engineering trick”^4^ that has brought pluripotent cells to the front stage as a tool for tissue replacement, disease modelling and animal engineering technologies^5^.

There are multiple pluripotent stem cell <u>types</u> that can be isolated from vertebrates, including rodents and human, typically annotated based on their donor cell-of-origin **(Fig. 1)**. Embryonic stem cells (ESCs) are isolated from the inner cell mass (ICM) of developing pre-implantation mouse or human blastocysts^6-8^. Epiblast stem cells (EpiSCs) are isolated from mouse postimplantation epiblasts^9,10^, however equivalent derivations haven’t been attempted with human embryos due to justified ethical complexities. Early rodent migrating PGCs can be converted *in vitro* into pluripotent ESC-like cells termed embryonic germ cells^11,12^. Mouse neonatal and adult spermatogonial stem cells can be reverted toward pluripotency and generate male germ stem cells (GSCs)^13-15^. The latter have the disadvantage of retaining only male imprint signature, which can increase tumorigenic potential^15^. Intriguingly, stable and validated EGs and GSCs have not been isolated from primates thus far^16,17^ **(Fig. 1)**.

**Figure 1.**
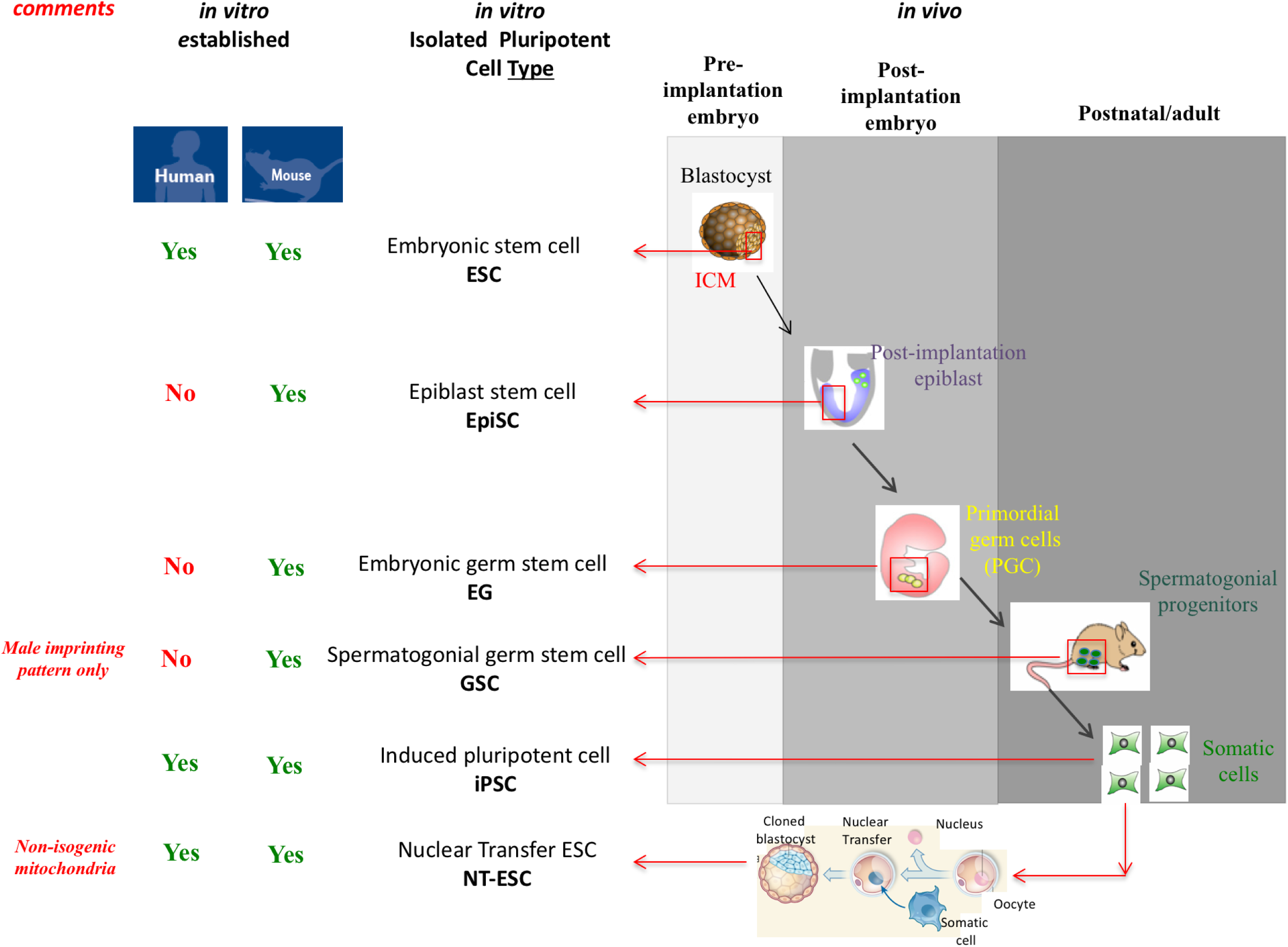
Routes for deriving different mouse and human pluripotent stem cell types. A variety of pluripotent cell types can be derived from different embryonic cells harvested at various stages of mouse or human development. Alternatively, somatic cells can be reprogrammed via somatic cell nuclear transfer (SCNT) or *in vitro* reprogramming via exogenous transcription factors to generate iPSCs. Post-implantation epiblast or germ cell lineage derived pluripotent cells (EpiSCs, EG and GSCs) have not been stably derived yet in human or other primates. For therapeutic purposes, iPSCs and NT-ESCs have the advantage of being genetically identical to donor original somatic cells. However, NT-ESCs will retain mitochondria of the a-nucleated donor female oocyte. Spermatogonial germ stem cells are generated from spermatogonial stem cells that have already established exclusive male/paternal imprinting pattern, and so do GSCs derived from them. If established from adult human males in the future, such maternal imprint-free characteristics will likely limit future therapeutic potential of human GSCs.

Somatic cell reprogramming provides alternative routes for isolating pluripotent cell types. Human and rodent somatic cells can be artificially reprogrammed into ESC-like cells following reprogramming via nuclear transfer, termed NT-ESCs^18-20^. Ten years ago, Yamanaka established direct *in vitro* reprogramming of somatic cells to pluripotency via ectopic expression of defined factors^21^, that yield induced pluripotent stem cells (iPSCs) without the need for oocytes or embryos^22–25^ **(Fig. 1)**. NT-ESCs and iPSCs offer the advantage of being able to generate patient specific pluripotent cells with nuclear DNA identical to the donor somatic cell, however mitochondrial DNA in NT-ESCs is non-isogenic and provided by the a-nucleated donor oocytes^26^. The latter can be an advantageous in settings aiming at correcting maternally inherited mitochondrial diseases^27-29^.

While the above overview pertains to classify different pluripotent cell <u>types</u> based on their tissue derivation source, the growth conditions used to expand such cells dictate the pluripotent <u>state</u> they attain (ICM-like, ESC-like, EpiSC-like state etc.)^30,31^. iPSCs generated in classical mouse ESC growth conditions yield ESC-like iPSCs; while those reprogrammed in EpiSC growth conditions yield EpiSC-like iPSCs^30,32^. The same analogy applies to explanting rodent ICM cells in ESC or EpiSC growth conditions^30,33^. In comparison to developmentally restricted mouse EpiSCs, ESCs are highly competent in generating high-contribution chimeric mice after microinjection into host-blastocysts, retain a pre x-inactivation state in female cell lines and reduced expression of lineage commitment factors^30,31^. Such attributes influence the utility of pluripotent cells in cell differentiation and animal transgenics. Thus it is of importance to understand and define different pluripotent states and configurations across different species^4^.

In this review we provide an integrative perspective on recent breakthroughs in understanding the diversity and complexity of pluripotent state regulation *in vitro.* This includes advances on preserving naïve pluripotency from non-rodent species and alternative pluripotent states. We highlight unresolved issues, key questions and future directions in this exciting front of stem cell research.

### Murine pluripotent states

Mouse ESCs were shown to reside in an ICM-like state^30^, referred to as the naïve state of pluripotency^31^, since they retain several molecular characteristics of ICM. A novel type of pluripotent cells, termed EpiSCs, was later derived from post-implantation rodent epiblasts^9,10^. In comparison to naïve ESCs, EpiSCs retain an alternative pluripotency configuration, referred to a primed pluripotency^31^. Profound molecular and functional differences exist between different pluripotent cells, which subsequently influence their characteristics, function and safety.

#### Growth conditions for naïve pluripotency

To grasp the biology of mouse naïve ESCs and their developmental context, it is of relevance to review the evolution of growth conditions devised to isolate such cells over the course of the last thirty years. Martin and Evans derived ESCs from 129-mouse strain^7,8^, by utilizing mitotically inactive embryonic fibroblast cells (MEFs) as feeder cells and foetal bovine serum (FBS)^34^. Leukaemia Inhibitory Factor (LIF) that activates the STAT3/JAK pathway was later identified as a key ingredient that allowed expansion of mouse ESCs in FBS/LIF conditions without MEFs^35,36^ **(Fig. 2)**. Such naïve mouse ESCs express hallmark pluripotency factors (e.g. Oct4, Nanog, Esrrb) and retain a pre-x inactivation state in female cell lines^37^ **(Fig. 3)**. Functionally, ESCs can populate the host pre-implantation mouse ICM upon microinjection into blastocysts, and generate high-contribution chimeras with colonization of the germ-line^37^.

**Figure 2.**
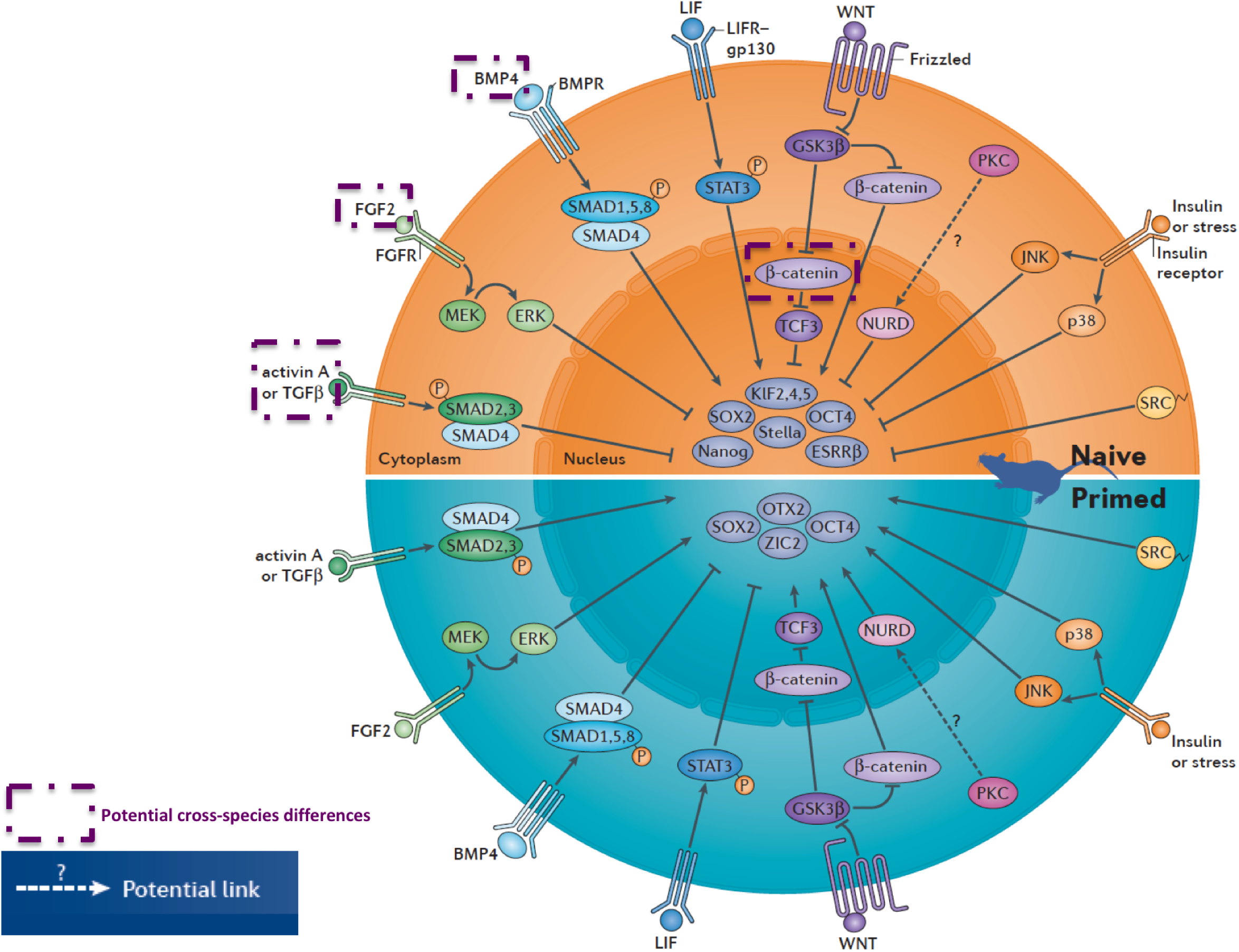
Signalling pathways and their influence on naïve and primed pluripotent states. Figure delineates different signalling pathways and their ability to positively or negatively regulate naïve and primed murine pluripotent cells. Please note how that the majority of signalling pathways delineated have opposing effects on murine naïve vs. primed pluripotent state stability (e.g. Lif/Stat3, Fgf2/Erk). It is important to highlight that other pathways not included in this scheme, are likely to be involved in such regulation and will likely be further characterized over the next years. This may include Hippo, Rho, Notch and NFkB signalling. Purple dotted boxes highlight signalling pathways that may be acting differently in mouse vs. human PSC regulation. More specifically, it remains to be fully understood whether low dose of TGFP, ACTIVIN/NODAL, nuclear PCatenin or FGF2 (MEK/ERK independent) signalling influence human naïve pluripotency differently than previously observed in rodent naïve ESCs. Dotted arrow indicates potential links that remain to be established. (Image is reused and modified from our Nature Reviews Molecular Cell Biology Poster (http://www.nature.com/nrm/posters/pluripotency/index.html)).

**Figure 3.**
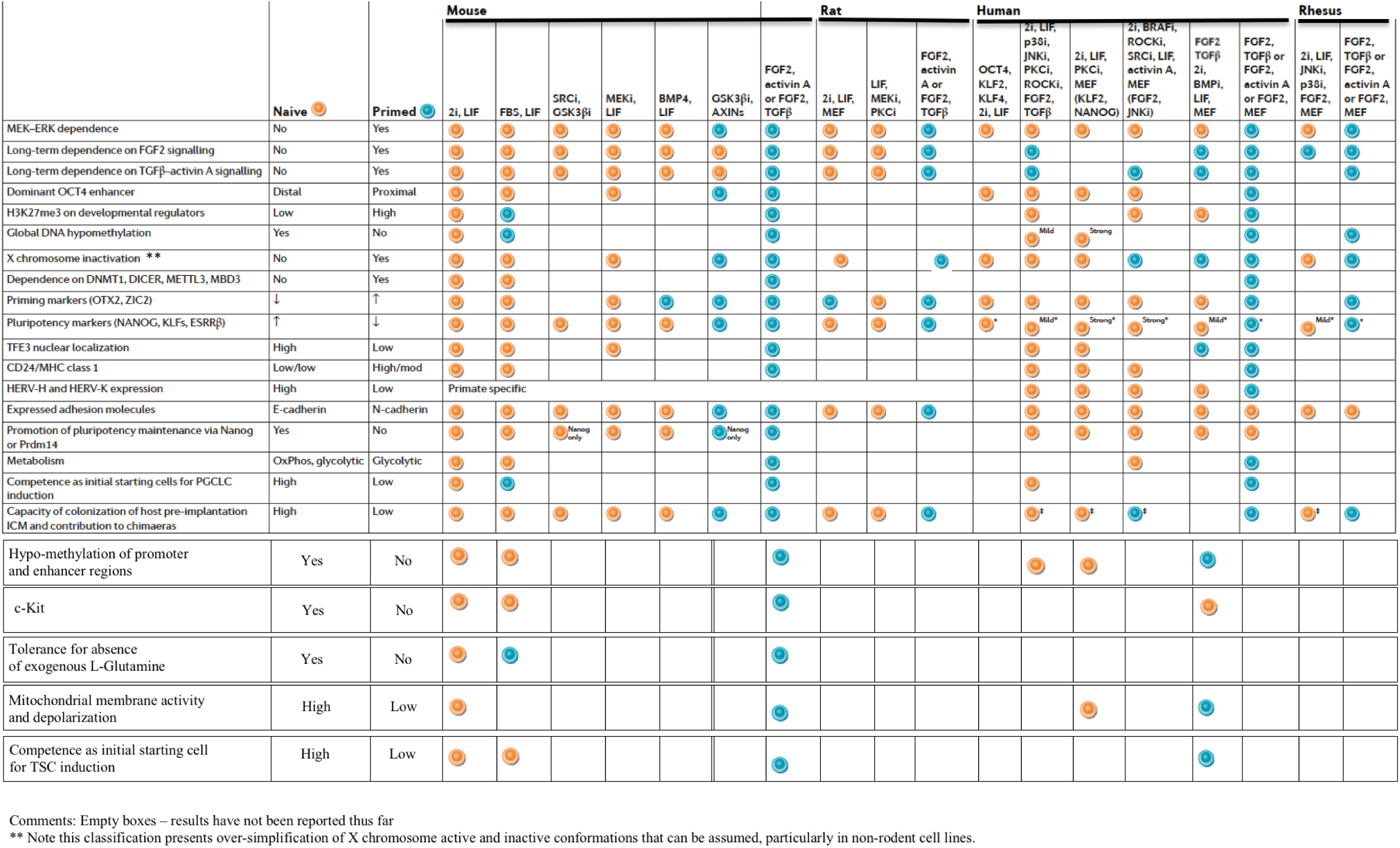
Naïve and primed pluripotent cell properties in different isolated PSCs. The scheme delineates in the first column on the left a list of different properties that distinguish between murine ESCs expanded in 2i/LIF (naïve) and murine EpiSCs expanded in Fgf2/Activin A (primed). The latter two states are used as a reference to annotate a variety of other naïve and primed conditions devised for mouse, rat, human or monkey PSCs. For each condition and for each stem cell property, we indicate whether it retains naïve-like (orange circle) or primed-like pattern (blue circle). Empty boxes indicate lack of characterization. This list of features is likely to expand with time and can be used to systematically annotate new pluripotent states isolated in unique conditions and from different species. (Image is reused and modified from our Nature Reviews Molecular Cell Biology Poster (http://www.nature.com/nrm/posters/pluripotency/index.html)).

Smith group described the first serum- and feeder-free defined conditions for expanding mouse ESCs by combining low-dose BMP4 with LIF^38^. Addition of small molecule inhibitors for MEK signaling increased ESC derivation efficiency and stability^39^. Developing different defined conditions, all involving MEK inhibitors, which can be used to isolate murine ESCs, extended the latter defined growth conditions. A cocktail combing 3 inhibitors termed ‘3i conditions’, was shown to stabilize pluripotent cells without LIF, indicating existence of redundant pathways to isolate ESCs *in vitro* that can compensate for lack of LIF/Stat3 signalling^40^. Notably, such cell configuration was labelled as “ground state pluripotency” since the cells in 3i were reported to grow independent of any exogenous signalling stimuli. However, this term is challenged by the fact that growth in these conditions heavily relies on GSK3 inhibition that mimics WNT stimulation; and on exogenous insulin that activates PI3K/AKT signalling^40^. Further, 2i/LIF conditions were adopted as an enhanced mean to expand murine ESCs, and the reduced proliferation in 3i conditions indicates a role for autocrine secreted FGF4 in promoting naïve ESC growth independent of Mek/Erk^30,41,42^ **(Fig. 2)**. Notably, complete genetic ablation of Erk1/2 is indispensable for maintaining rodent naïve ESC survival and does not yield an identical phenotype to Mek inhibition, suggesting additional roles for Mek inhibition in maintaining ESCs stability that are Erk independent^43^.

“Alternative 2i” conditions, involving small molecule inhibitors for Gsk3 and Src pathways, yield germ-line competent ESCs^44^ **(Fig. 2)**. Atypical PKC small molecule inhibitor Go6983 (aPKCi) was identified as another stimulator for isolating murine ESCs together with LIF and/or MEK inhibitors^45^. Single cell RNA-seq analysis has proven equivalent global heterogeneity between different naïve conditions, however the difference exists in the identity of genes that underlie heterogeneity in each condition^46^.

Enriched conditions were important for deriving ESCs from other mouse strains that have until recently been considered “non-permissive” for deriving naïve ESCs. While 129-mouse strain derived ESCs could be expanded in FBS/LIF conditions, for other mouse strains like non-obese-diabetic mice (NOD), supplementation of 2i or GSK3 inhibitors is essential for both derivation and maintenance^30,42^. 3i and 2i/LIF conditions yielded rat ESCs, however these conditions are suboptimal^47,48^. LIF/MEKi/aPKCi constitute a more robust method to support rat ESCs^49^ **(Fig. 2,3)**.

The above findings underscore the relevance of analysing rodent ESCs expanded in different naïve conditions and from different genetic backgrounds. Further they emphasize the importance of other signalling pathways that remain to be deeply characterized in the context of pluripotency **(Fig. 2)**. SRC functions as a downstream target of MEK/ERK and Calcineurin-NFAT signalling to promote ESC differentiation^50^, and its inhibition promotes naïve pluripotency^44,50^. NF-kB inhibition has been implicated as a downstream effector underlying aPKC inhibition mediated support of naïve pluripotency^45^, however other pathways, like Mbd3/NuRD repressor complex, can be neutralized by aPKCi (unpublished observations, J.H.H).

Hippo signalling pathway regulates epiblast vs. trophoblast segregation in late mouse morulas, and is highly active in pluripotent epiblast cells, leading to exclusion of Yap/Taz effectors from the nucleus^51,52^. Piccolo and colleagues indicated that Yap/Taz depletion in mouse naïve ESCs expanded in 2i/LIF further enhances their resistance to differentiation^52^, while Lian et al. have indicated Yap/Taz as essential regulators for stability of naïve ESCs expanded in FBS/LIF^53^. Reanalysis of these findings in different conditions might resolve these seemingly opposing results **(Fig. 3)**.

It should be noted that signalling pathways are often pleiotropic and may simultaneously have positive and negative effects on naïve pluripotency. For instance, nuclear β-catenin stabilization following Gsk3 inhibition, promotes naïve pluripotency via neutralizing the repressive activity of Tcf3 on its bound target genes in the nucleus^54^. Cytoplasmic β-catenin promotes naïve pluripotency via increasing E-cadherin membrane stability^55^ **(Fig. 2)**. However, nuclear β-catenin can induce mesodermal gene expression through its Lef co-effectors^47^. Yet, such differentiation priming effects are outweighed by naïve pluripotency promoting functions of β-catenin under optimized conditions. LIF has also been shown to promote primitive endoderm specification in naïve pluripotency growth conditions^56^. Such “non-purist” effects should be kept in mind when dissecting the role of signalling pathways on pluripotency.

The ample conditions to grow naïve murine ESCs have been important for better understanding and revisiting the roles of several classical pluripotency regulators. While Nanog was first purported to be absolutely essential and irreplaceable for establishing naive pluripotency through iPSC reprogramming, cell fusion or EpiSC reversion^57^, multiple conditions enable reprogramming of Nanog null donor cells *in vitro*^58-60^. Still however, Nanog null ESCs cannot be derived from mouse ICMs, indicating that while Nanog is indispensible for establishing pluripotency *in vivo,* it is dispensable during *in vitro* induction^61^. The latter example highlights that *in vitro* pluripotency maintenance cannot be considered “authentic”, as some *in vitro* conditions can potentiate the robustness of the naïve pluripotency program and compensate for deficiencies that are not sustainable *in vivo.* Similarly, Klf2 knockout embryos do not present lethality at the pre-implantation stage, and naïve ESC in 2i/LIF or FBS/LIF conditions can tolerate Klf2 ablation^62^. However ESCs in 2i only conditions, cannot sustain loss of Klf2^62^, as LIF can compensate for the lack of Klf2.

Another emerging regulatory principle is that not all factors expressed in the ICM or ESCs necessarily promote naïve pluripotency, and some of them in fact are promoting its dissolution. However, they are tolerated by ESCs *in vitro* due to the optimized and enriched *in vitro* growth conditions used. For instance, Tcf3 binding represses the expression of its naïve pluripotency promoting target genes, leading to their partial repression but is tolerated in FBS/LIF naïve conditions. However, Tcf3 neutralization by adding GSK3i boosts naïve pluripotency^54^. In a similar manner, mouse ESCs tolerate Mbd3/NuRD complex expression although it partially represses naïve pluripotency targets^63^. However, genetic ablation of Mbd3 leads to upregulation of master regulators of naive pluripotency and allows LIF independent growth^63^. Consistently, derivation of Mbd3 KO ESCs from ICMs is uncompromised in 2i/LIF conditions^64^. In summary, both Tcf3 and Mbd3 are expressed in the ICM and ESCs, likely to set the stage for terminating naïve pluripotency. Thus, the molecular characteristics of a pluripotent state fixated in a certain condition represent the net outcome of conflicting stabilizing and destabilizing factors simultaneously residing and conflicting in that state^65^.

Collectively, it is important when analysing function of pluripotency regulators to systematically compare different naïve growth conditions, genetic backgrounds and *in vivo* context^66^. Such integrative analysis will likely unravel additional layers of underappreciated complexity and may resolve some conflicting results^52,53^.

#### Growth conditions for primed pluripotency

Primed EpiSCs were derived from post-implantation epiblasts of rodents in Fgf2/Activin A conditions^9,10^ **(Fig. 1)**. EpiSCs are capable of differentiating into cells of all three germ layers *in vitro* or in teratoma assay, and thus are pluripotent. However, they are inefficient in yielding chimeric animals once injected in pre-implantation epiblasts **(Fig. 3)**, likely because they have altered molecular characteristics and correspond to a more advanced developmental stage in comparison to the host pre-implantation environment^9,10^. They can however form low contribution chimeric embryos, when injected into host-post implantation embryos *ex-vivo^67^.*

While EpiSCs maintain Oct4 and Sox2 expression, they down-regulate most of the other pluripotency factors including Nanog, Esrrb, Klf2 and Klf4^3^. EpiSCs have not undergone differentiation, but they upregulate lineage commitment factors like Otx2, Brachyury and Zic2^68^. Epigenetically, EpiSCs retain distinct characteristics from naïve ESCs: they inactivate X chromosome in females, upregulate global DNA methylation levels and acquire H3K27me3 at developmental regulators^69,70^. Enhancer landscape is rewired between naive and primed states^68^, and developmental regulator gene associated “seed enhancers” convert from a dormant to an active state in EpiSCs, thus pre-marking lineage differentiation bias of primed PSCs^71^. Summary of divergent signalling and molecular characteristics between murine primed and naïve cells are highlighted in **Fig. 2-3**.

At the regulatory level, naïve and primed pluripotent cells have been shown to retain opposing dependence on epigenetic repressors^66^ **(Fig. 4)**. Naïve ESCs expanded in FBS/LIF or 2i/LIF tolerate loss of epigenetic repressors like Dnmt1, Dicer, Eed, Mbd3 and Mettl3^66^, and renders these cells “hyper–naïve” and resistant to differentiation^66,72^ **(Fig. 4)**. On the contrary, murine primed pluripotency maintenance and viability depends on these regulators, and their ablation destabilizes the murine primed pluripotent state^66^ **(Fig. 4)**. Defining how the depletion of each of these repressors precisely destabilizes the primed configuration is of future interest.

**Figure 4.**
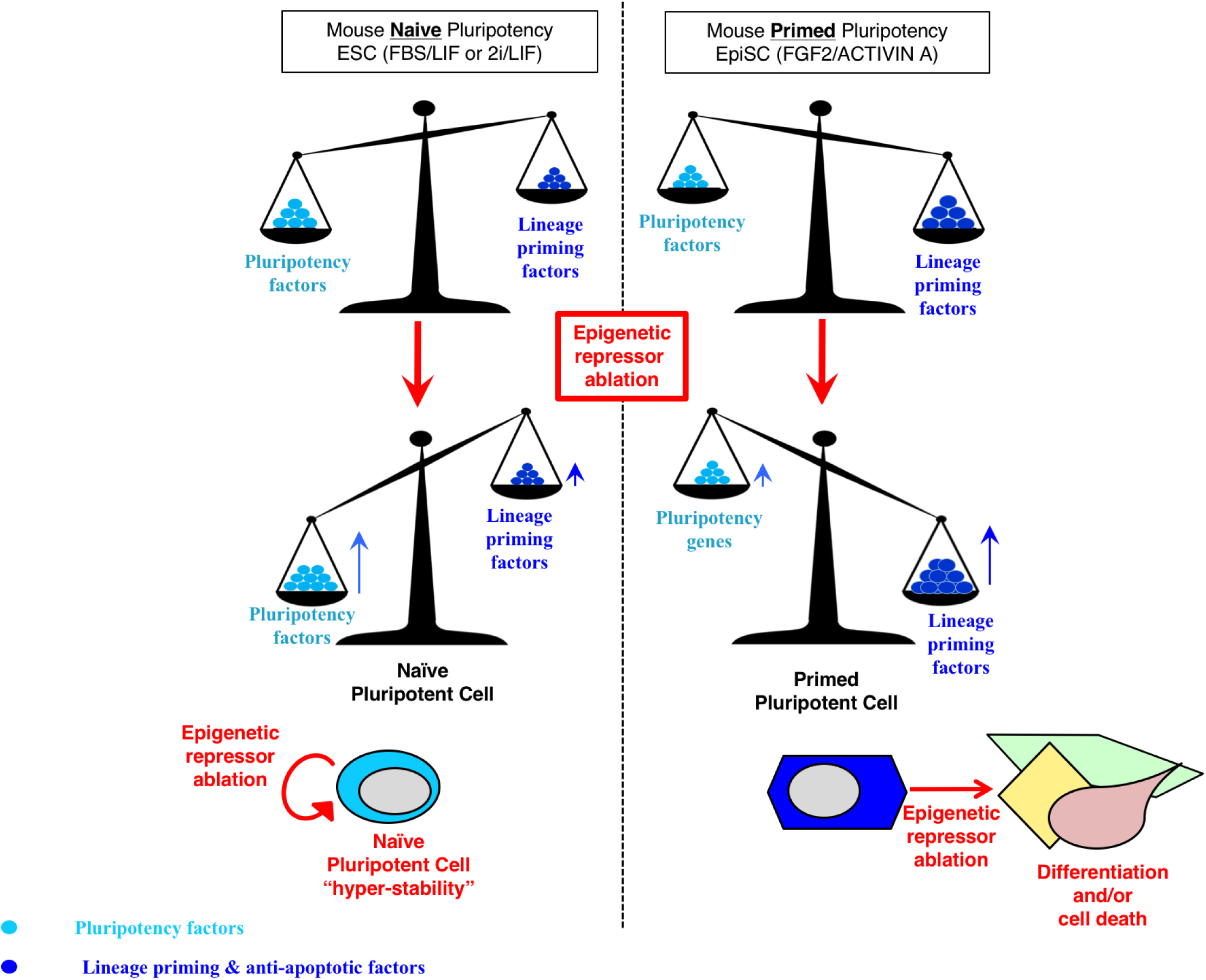
Opposing influence for epigenetic repressors on murine naïve and primed pluripotent cells. Murine naïve and primed pluripotent cells do not only differ in their dependence on distinct signalling pathways or in their epigenetic profile, but also in their lineage decision-making. Murine naïve ESCs expanded either in 2i/LIF and FBS/LIF conditions, tolerate complete loss of epigenetic repressors such as Dnmt1, Mbd3, Dicer, Dgcr8, Mettl3, Eed, Ezh2. Further, the latter modifications strengthen the equilibrium in favour of pluripotency promoting factors and generates “hyper-naïve” pluripotent cells that are relatively more resistant to differentiation and can tolerate withdrawal of LIF cytokine. Primed EpiSCs generally respond in an opposite manner to the complete ablation of such repressors. Established murine primed EpiSCs naturally down regulate pluripotency factors and upregulate lineage priming factors. Ablation of repressors at this stage tips the balance toward differentiation and/or compromises cell survival. It should be noted that ablation of different repressors (e.g. Mettl3 vs. Dgcr8) overall negatively influences primed pluripotency stability, this does not necessarily mean that the downstream events leading to the state collapse are identical, and thus should be thoroughly dissected *in vitro* and *in vivo.*

Alternative growth conditions to expand murine EpiSCs have begun to emerge. Ying and colleagues showed that simultaneous use of a GSK3 inhibitor (that induces β-catenin stabilization), together with a Tankyrase small molecule inhibitor IWR1 (that up regulates Axin1/2 levels thus leading to retention of β-catenin in the cytoplasm) maintain novel primed EpiSCs without exogenous Fgf2/Activin A supplementation^73^ **(Fig. 3)**. Removal of IWR1 leads to increased nuclear β-catenin shuttling and EpiSC differentiation^73^. The mechanisms by which cytoplasmic β-catenin prevents EpiSC differentiation remains to be uncovered^73^. It is tempting to speculate whether the recently described ability of cytoplasmic APC/Axin/β-catenin destruction complex to act as a sequestration “sink” for Yap/Taz and prevent their nuclear shuttling^52^, is involved in the ability of GSK3i/IWR1 conditions to maintain EpiSCs. Notably, the latter alternative EpiSC state is different from EpiSCs expanded in classical Fgf2/Activin A conditions, and retains higher expression of naïve markers^73^, and are thus relatively “less primed” **(Fig. 3)**.

Recent studies indicate that different primed conditions can endow EpiSCs with region specific characteristics of the post-implantation epiblasts. EpiSCs expanded in Fgf2/Activin A correspond transcriptionally and functionally to anterior late-gastrula primitive streak cells^74^. Alternative FGF2/IWR1 conditions generate murine EpiSCs corresponding rather to posterior-proximal epiblasts^75^. Further, even in classical Fgf2/Activin A conditions distinct subpopulation of EpiSCs can co-exist, each representing different stages of post-implantation embryonic development^76^.

Finally, the time length at which pluripotent cells are maintained under primed conditions greatly influences their characteristics and functionality^77^. Counter intuitively; while murine PGCs are specified from the post-implantation epiblast *in vivo,* EpiSCs maintained *in vitro* for more than 7 days in FGF2/Activin A, lose competence to generate PGCs in response to Bmp4^77^. Starting with naïve cells and inducing brief priming for 2-4 days, yields distinct primed cells highly competent for generating PGCLCs, termed EpiLCs^77^. The latter are transcriptionally more similar to *in vivo* post-implantation epiblast than EpiSCs^77^. Thus, the above paradigm indicates another aspect of artificial features that can be acquired by pluripotent cells once expanded indefinitely *in vitro,* in contrast to them *in vivo* “counterparts” that transiently exist during development.

Studies involving clonal lines and single cell analysis will provide deeper understanding of features of region specific EpiSCs and shortly after *in vitro* induction from a naïve state in different priming conditions^74^. This may help understand how lineage priming is established at the single cell level during these key early developmental transitions^74,75^ and might be relevant for optimizing other differentiation protocols and predicting PSC behaviour.

#### Interconversion between naïve and primed states

As somatic cells can be reprogrammed into a naïve ESC-like state via combined overexpression of pluripotency factors together with LIF, primed EpiSCs can also be reverted to naive iPSCs. Overexpression of Klf4 or Myc in EpiSCs, under LIF containing conditions, generates naïve ESCs^30,78^. FBS/LIF signalling alone can be sufficient to induce such conversion from permissive mouse genetic backgrounds (i.e. 129 strains)^79^, but not from “non-permissive” ones like NOD, where supplementation of small molecules like 2i is necessary^30^. Other factors like Nanog, Prdm14 and Esrrb have been shown to synergistically induce and boost the efficiency of this process^80,81^. Explanting post-implantation E5.5-E7.5 epiblasts in naïve conditions also reverts them into naïve PSCs^30,79^. The opposite conversion is attainable from *in vitro* and *in vivo* isolated naïve cells, as expanding murine naïve PSCs or ICMs in primed conditions leads them to gradually adapt an EpiSC state^30,33,78^.

Studies focusing on *in vitro* molecular changes accompanying naïve to primed pluripotency conversion have unravelled key events in understanding mechanisms of reprogramming^69^. Naïve ESCs expanded in 2i/LIF retain global hypo-methylated levels in both promoters and gene bodies, highly similar to those measured in ICMs^82,83^. When transferred into LIF/FBS naïve conditions, this is accompanied by an increase in global DNA methylation levels, however promoter and enhancer regulatory regions remain protected from invasion by DNA methylation^84^. Only after transfer into primed Fgf2/Activin A EpiSC inducing conditions, DNA methylation accumulates over enhancer and promoter regulatory elements^84^.

Transitioning naïve FBS/LIF PSCs into 2i/LIF conditions leads initially to changes in Oct4, Sox2 and Nanog pluripotency factor occupancy^85^. Changes in H3K27me3 deposition and enhancer landscape follows later, likely in response to the rewiring in transcription factor binding^85^. Downregulation in DNA methylation follows next, which has mainly been attributed to downregulation in *de novo* DNA methyltransferase enzymes^82^. It should be noted however, that ablation of Dnmt3a/b in ESCs in FBS/LIF condition does not lead to such rapid loss of DNA methylation^86^, and other yet to be identified events might be involved in this rapid 2i induced epigenetic response. MEK/ERK inhibition influences polycomb interactions and leads to decreased occupancy by PRC2 and decreased phosphorylation on the CTD of PolII on lineage commitment genes^87^, leading to loss of H3K27me3 and increased PolII pausing at bivalent developmental regulatory loci^69^. Analysing other defined naïve pluripotency growth conditions (2i/LIF/PKCi, alternative 2i etc.) and in other rodents, will be important for discerning redundancies and specificities of different singling pathways and how they cross-talk with chromatin organization.

#### Human conventional pluripotent cells

Thomson group first isolated human ESCs from bastocysts^6^. Surprisingly, they were drastically different from murine ESCs in their characteristics and tissue culture requirements. FGF2 and TGFβ1/ACTIVIN, but not LIF, signalling are at the core signalling modules maintaining such conventional human ESCs derived from the ICM, or iPSCs obtained via direct *in vitro* 88 reprogramming.

#### A primed pluripotent state

Differences between conventional human and mouse ESCs had been initially attributed only to unknown species genetic differences, since human ESCs were also derived from the ICM and not from post-implantation stages. However studies on different mouse strain derived stem cells have discerned a scenario where ICM cells can adapt *in vitro* into a primed state if naïve conditions that match the requirements of the particular genetic background of donor embryos used, are not devised^30^. Specifically, NOD mice are relatively “less-permissive” than 129 mice to yield naïve ESCs/iPSCs, as LIF alone is not sufficient to maintain NOD naïve pluripotency and 2i/LIF are permanently required to stabilize and maintain this state *in vitro* in NOD PSCs^50^ Further, ICMs from both 129 and NOD strain expanded in primed conditions yield EpiSCs that are indistinguishable from EpiSCs derived from E6.5 embryos^30^ or *in vitro* from already established ESCs^30^.

The relevance of the latter *in vitro* priming scenario to dictating conventional human ESC identity is supported by the fact that human conventional ESCs/iPSCs retain a great milieu of primed pluripotency features. This includes low expression of naïve pluripotency markers (e.g. TFCP2L1, DPPA3), deposition of H3K27me3 over developmental genes, lack of exclusive nuclear localization of TFE3, loss of pluripotency upon inhibition of MEK/ERK pathway, lack of global hypomethylation as seen in ICM cells, lack of a pre x-inactivation state in most conventional female PSCs lines^70,89,90^. Further, human primed ESCs do not tolerate complete loss of DNMT1^86^, similarly to what has been shown for mouse EpiSCs^66^ **(Fig. 4)**. Complete human KO ESCs have not been obtained thus far for DICER, MBD3 or METTL3^64,66^ **(Fig. 3)**.

#### Less primed than murine EpiSCs

In spite of the above, it is of importance to realize that human conventional/primed ESCs are not identical to murine EpiSCs, and can be considered relatively “less primed”. For instance, human ESCs do not upregulate FGF5 and N-CADHERIN as seen in murine EpiSCs, and express high-levels of E-CADHERIN as detected in mouse naïve ESCs^70^. Human ESCs express high levels of some naïve markers like NANOG, PRDM14 and REX1 that are residually expressed in murine EpiSCs^91^. Moreover, human primed ESCs are functionally dependent on NANOG and PRDM14 and their ablation induces differentiation^91^. DNA methylation distribution in human ESCs shows they rather correspond to murine naïve ESCs expanded in FBS/LIF conditions, rather than mouse FGF2/Activin A expanded mouse EpiSCs, as their promoters are protected from invasion by repressive DNA methylation^84,92^. Further, while murine EpiSCs demonstrate exclusive TFE3 cytoplasmic localization and naïve 2i/LIF ESCs show exclusive nuclear TFE3 localization^93^, human primed ESCs show an intermediate configuration where TFE3 is present in both the cytoplasm and the nucleus^70^.

### Human naive pluripotent cells

The metastability of naïve and primed pluripotent state depending on the growth conditions applied^30^, and the stringency in requirement for exogenous naïve pluripotency promoting factors to isolated naïve PSCs from previously “non-permissive” rodent strains^30,31^, have underscored a scenario of whether unique and more stringent conditions can be applied to isolated previously unidentified alternative naïve-like pluripotent states in humans.

#### Transgene-dependent generation

2i/LIF conditions are not sufficient to maintain naïve human ESCs or iPSCs^94^. However, additional transgene expression can induce an artificial transgene dependent state that may be of considerable interest. Continued exogenous OCT4/KLF4 or KLF2/KLF4 transgene expression can maintain human ESCs/iPSCs in a unique pluripotent state in 2i/LIF conditions^94^. Recently, these observations were extended by optimizing over-expression of KLF2/NANOG transgenes, allowing expansion of human naïve iPSCs in 2i/LIF^95^. Smith and colleagues overexpressed KLF2 and NANOG transgenes in primed ESCs and expanded them in 2i/LIF/aPKCi conditions ^96^. These cells exhibited extensive DNA hypomethylation, and strong upregulation of naïve markers like TFCP2L1, KLF2, and KLF4. However, as KLF2 is not expressed in the human ICM^97^, and as 2i/LIF/aPKCi are insufficient to convert primed ESCs without exogenous transgene induction^96^, and as transgene-free cells remain to be validated in 2i/LIF/aPKCi conditions, it is unclear whether this state is indefinitely stable without retaining possibly leaky transgenes or MEFs. Further, independent examination of DNA methylation landscape in these cells indicates aberrant global loss of imprinting, excessive hypomethylation of endogenous retroviral genes^89,98^. Finally, while 2i/LIF/aPKCi conditions do not contain exogenous FGF or TGFβ1/Activin A cytokines, applying only short-term inhibition of FGFR/TGFR signalling is not sufficient evidence to validate FGF/TGF/Activin A signalling independence^96^ **(Fig. 2)**. Indeed, unlike in mice, human pluripotent ICM collapses upon treating blastocysts with small molecule inhibitors for TGF/ACTIVIN signalling^97^.

While the field has shifted to study transgene independent conditions as detailed below, it should be noted that such transgene dependent states might be important, since it may be possible that robust naïve pluripotency currently obtained in mouse ESCs is a rodent specific phenomenon. Capturing human naive PSCs identical to those obtained from mice might still involve genetic modifications. Nevertheless, the latter studies provide evidence for the possibility to generate alternative pluripotent states in humans and other species^30,94^.

#### Transgene independent generation

Our team was the first to describe naïve conditions, designated as NHSM (Naïve Human Stem cell Medium), which entail complete ablation of MEK/ERK signalling and are compatible with indefinitely expanding genetically unmodified human PSCs both in MEF-containing and -free conditions^70^. These naïve MEK independent pluripotent cell lines could be derived from human pre-implantation embryos, through *de novo* iPSC generation, or from previously established primed ESCs/iPSCs^70^. NHSM conditions contain 2i/LIF together with P38i, JNKi, aPKCi, ROCKi, low doses of FGF2 and TGFβ1 (or Activin A), and render human PSCs more similar, but not identical, to murine naïve PSCs^70^ (Fig. 2,3). In fact, these cells have features of so called naïve 2i/LIF “ground state” pluripotency, which are not found even in naïve mouse ESCs expanded in FBS/LIF. This includes exclusive nuclear localization of TFE3, and cleansing of H3K27me3 over developmental genes^69,70,93^. Transcriptionally, these cells down regulated expression of lineage commitment markers like OTX2, ZIC2 and CD24 and moderately upregulated pluripotency genes (more prominently on MEFs and when aPKCi is used)^70,99^. Enhancer rewiring has been attained in these human naive PSCs as seen with mouse cells^71^. The cells exhibited downregulation in DNMT3B^100^ and a mild global decrease in global DNA methylation levels, while maintaining imprinting integrity and chromosomal stability^70^. While conducting chimeric analysis with human PSCs and using human embryos as hosts is ethically and legally forbidden, these human naïve PSCs cells showed better integration into blastocysts upon microinjection into host mouse morulas and were able to contribute at low-grade levels in mouse embryos up to E10.5-E17.5^70^ **(Fig. 3)**.

Important publications describing alternative conditions that yield human MEK independent naïve pluripotent cells emerged after, each producing cells with different enhanced molecular properties **(Fig. 3)**. A combination of 2i/LIF, ROCKi, BMPRi, high doses of FGF2 and TGFB1 were able to maintain human PSCs only in the presence of MEFs^101^. These cells demonstrated transcriptional upregulation of pluripotency markers like STELLA and KLF5. Jaenisch team described conditions^95^ that adopted most of components found in NHSM^70^ (i.e. 2i/LIF, ROCKi, Activin A (instead of TGFβ1) - with or without FGF2 and JNKi) and supplemented inhibitors for BRAF and SRC pathways (conditions termed 5i/LA-MEF with optional inclusion of JNKi or FGF2). In comparison to previous studies, cells in 5i/LA-MEF conditions demonstrated a more impressive upregulation of naïve pluripotency markers. However, the cells did not down regulate DNMT3B, maintained an inactive X chromosome state in female cell lines, and demonstrated an unusual pre-ICM signature^95^. Intriguingly, the process of converting primed cells back to naïve state in 5i/LA-MEF conditions is inefficient, taking two weeks to isolate initial clones that retain a slow growth rate^95^. Further, these conditions exclusively yield chromosomally abnormal cell lines^95^. Thus, it remains to be determined whether such chromosomal abnormalities are in fact inherent to 5i/LA-MEF cells and dictate the properties described for this state^95^, and are being selected for during this inefficient conversion process. Finally, DNA methylation profiling and whether these cells maintain epigenetic imprinting integrity is another important aspect that remains to be evaluated.

It is clear from the above summaries that none of the many already published conditions generate human naïve PSCs that are identical to mouse ESCs or human icm^97,102,103^. However, these studies implicate new signalling pathways and pave avenues for further optimization and characterization of such novel PSCs **(Fig. 2)**. Mechanistically it will be interesting to test whether there is a connection between RAF and aPKC inhibition and the influence of modulating WNT signalling by applying GSK3i/IWR1 module^73^ on human naïve PSCs. Further, RHO signalling has been shown to promote YAP/TAZ nuclear localization in primed human ESCs and sustain their pluripotency^104^. Thus it remains to be defined whether ROCKi influencing naïve pluripotency characteristics^70^ via YAP/TAZ modulation.

The role of MEK independent FGF2 and TGFβ1/ACTIVIN A signalling, either autocrine or exogenously provided at low doses, remains to be understood in human naïve PSCs. The latter demonstrate upregulation of Activin like ligand GDF3^96^, and human, but not mouse, ICM cells abundantly express Activin receptors^97^. Thus, it is tempting to speculate that primed human ESCs are relatively less primed than murine EpiSC due to differences in response to Activin-like ligands, where it might promote some naïve features in human^70,95^, but not in mouse. To conclude, systematic analysis of the response of pluripotent states from different species to a variety of TGF ligand family members is of importance **(Fig. 2)**, while the possibility to generate human PSCs that are entirely independent of FGF and/or TGF signalling cannot be excluded.

### Differences between mouse and human epiblasts

Recent studies focusing on single cell RNA-seq of human pre-implantation embryos are starting to provide answers to some of the questions highlighted above. While human and mouse blastocyst do not display morphological differences, they retain profound molecular differences at the cellular level^97^ **(Fig. 5a,b)**. Human ICM epiblast cells do not express genes that are considered important pluripotency factors in mouse, such as KLF2 and ESRRB. Instead, KLF17 might have a human-specific role in the ICM^97^. ERAS, an ESC specific form of RAS, became a pseudo-gene in humans^105^, and Eras null mouse ESCs propagate slowly in FBS/LIF conditions^105^. Non-human primate (Marmoset) pluripotent epiblast is similar transcriptionally to human, and very different than mouse^106^. This includes lack of KLF2, NR0B1 and BMP4 transcription, while acquiring high expression of NODAL and its downstream signalling mediators in human and marmoset naïve ICM^106^.

**Figure 5.**
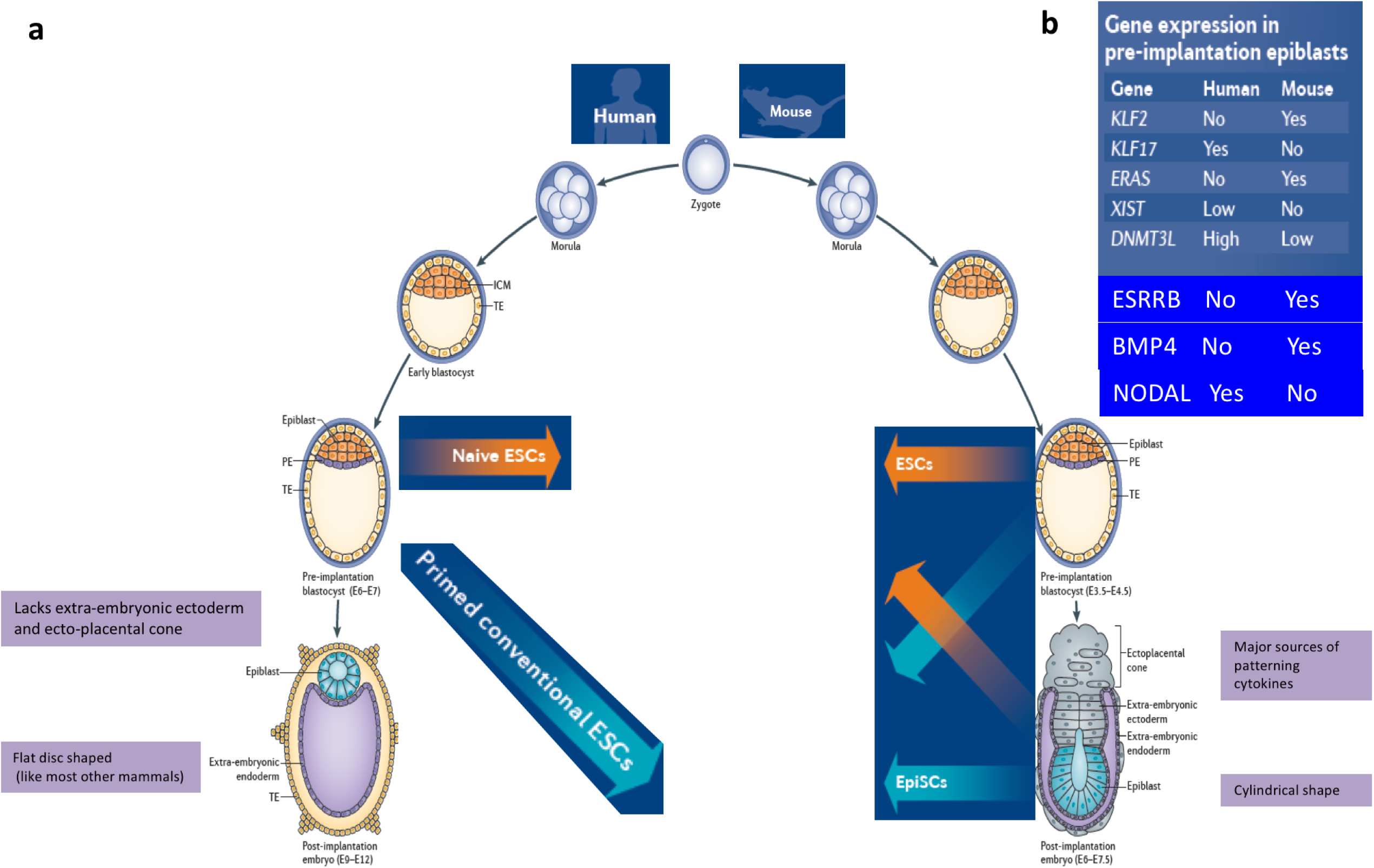
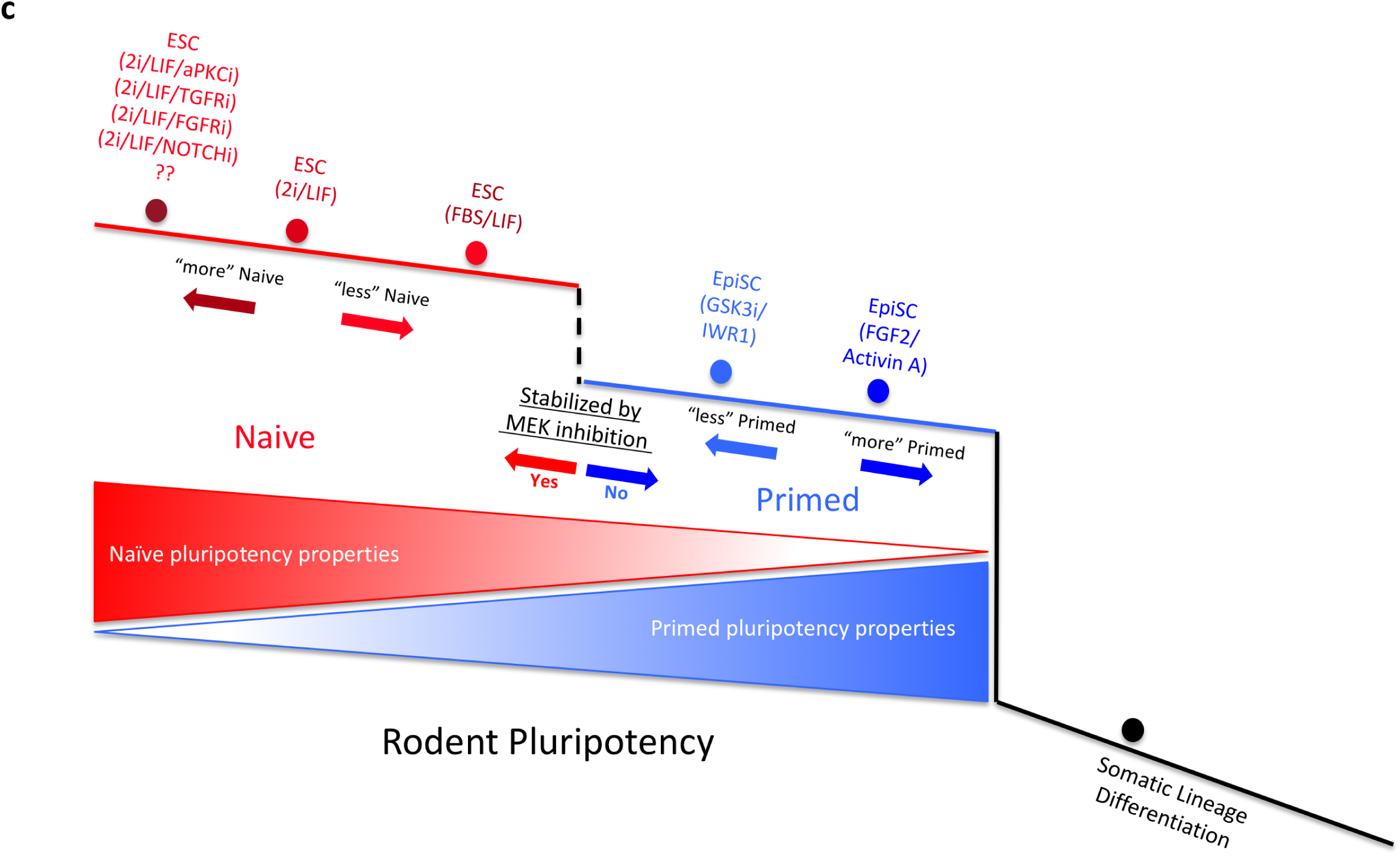
Key differences between mouse and human pre and post-implantation development *in vivo.* (a) Scheme delineates similarities and differences in mouse and human early pre- and post-implantation development. While mouse and human embryos are not morphologically different up to the blastocyst stage, there are striking transcriptional differences as summarized in (b). Further, at the postimplantation stage, morphological differences in human vs. mouse embryo shape become striking including (i) differences in epiblast shape and (ii) extra-embryonic structures, as delineated in (a). In mouse, naïve ESCs (orange arrows) can be derived from ICM or post-implantation epiblast when naïve growth conditions are applied. Primed EpiSCs can be derived from post-implantation epiblast or ICM when primed culture conditions are used. Thus, growth condition rather than source dictates pluripotent state configuration acquired *in vitro.* A similar scenario applies for derivation of naïve and primed cells from human ICM depending on the growth conditions used. PSC derivations or molecular analysis on human post-implantation embryos cannot be conducted due to ethical issues. PE-Primitive Endoderm; TE – Trophectoderm; ICM-Inner Cell Mass. (c) A model to explain the relativity of naivety within the naive to primed pluripotency spectrum. One major molecular and functional criterion that can be considered for separating naïve and primed pluripotent cells is their ability to maintain and stabilize their pluripotent state upon induced blocking of MEK activity (dashed black line). Within the naïve or primed pluripotent states, it is difficult to describe the pluripotent state of the cells in absolute terms, as naïve cells can have, to some extent, primed pluripotency features. Similarly, within the primed pluripotency spectrum, primed PSCs expanded in different conditions have different features and varying degrees of naivety (**Figure 3**). Finally, it is possible that supplementation of 2i/LIF conditions with small molecules such as aPKCi, FGFRi or NOTCHi will further consolidate naïve pluripotency features, particularly from other rodents like rats whose stability in 2i/LIF feeder free conditions should be further improved. Full annotation of different human pluripotent states will allow charting an equivalent landscape for human and monkey PSCs. (Images in **a-b** are reused and modified from our Nature Reviews Molecular Cell Biology Poster (http://www.nature.com/nrm/posters/pluripotency/index.html). Image in c is reused and modified from our Figure 3 in Manor et al. Curr. Opin. in Genetics & Development).

At the post implantation stage, major differences exist between rodent and human embryos **(Fig. 5a,b)**. Rodents are rather unusual as their post-implantation epiblast assumes egg-like cylinder shape, while in humans the post-implantation epiblast assumes a flat disc shape, like in most other mammals. While it might be impossible to conduct single cell analysis on early human post-implantation epiblasts, non-human primates might provide some relevant insights. Collectively, these species differences might directly influence the distinct pluripotent characteristics observed in PSCs from different species *in vitro* and their distinct growth requirements. Further, they are of relevance for understanding the developmental context of human *in vitro* isolated pluripotent cells.

### A framework for classification of pluripotent states

The advent of different conditions to isolate human naïve PSCs with distinct characteristics, and the limitations in conducting chimeric analysis in humans, simulate discussions on classifying pluripotent states. It is often claimed that ability to derive ESCs for human ICMs in a newly devised growth condition constitutes “a gold standard” for proving naivety. However, it should be kept in mind that the pluripotent state identity is eventually dictated by the derivation growth condition and not by whether their source was from the pre- or post-implantation epiblast^30,33^, or PGCs^82^. Utilization of OCT4 distal vs. proximal enhancer element as a binary distinguishing marker can be also misinterpreted^95^. Both OCT4 distal and proximal enhancer elements are active in naïve and primed states, both in humans and mice^107,108^. The difference rather emerges from their relative activity levels (high/low) and dominance.

Relying on a single attribute marker or functional test is limiting and must be accompanied by systematic analysis of ever increasing characteristics that continue to be uncovered for different pluripotent states **(Fig. 3,4)**. Nevertheless, in our opinion, a molecular and functional characteristic that can be considered as a major divider between naïve vs. primed pluripotent states is their response to the challenge of blocking MEK signalling **(Fig. 5c)**. Human conventional ESCs and mouse EpiSCs rapidly collapse upon MEK inhibition, while naïve pluripotent cells rather tolerate and consolidate their naivety following this challenge^94^. Emphasizing this functional quality is supported by the ability of MER/ERK inhibition to expand murine epiblast in ICMs and that it signifies consolidation of naïve pluripotency *in vivo*^109^.

Within the domains of the naïve and primed ground states of pluripotency, it is clear that if one considers a list of many naïve and primed pluripotency features originally described for mouse naive 2i/LIF ESCs and primed FGF2/Activin A EpiSCs, different pluripotency growth conditions can simultaneously endow a mix of primed and naïve properties in the same cell type **(Fig. 3)**. As such, pluripotent states can be classified as “more naïve” or “more primed” by having more of such properties **(Fig. 5c)**. Human primed ESCs have a number of naïve pluripotency features (e.g. protection of promoter regions from hypermethylation, dependence on Nanog), and recent comparative analysis with single cell RNA sequencing of human blastocysts suggests that some conventional human ESC lines maybe transcriptionally relatively less primed than previously thought^97^. Murine naïve ESCs expanded in FBS/LIF conditions can give rise to “all-ESC” chimeric embryos and tolerate Dnmt1/Mettl3 ablation, however they are globally hypermethylated and acquire H3K27me3 over developmental genes as seen in EpiSCs (Fig. 3,5c). Thus, the latter conditions reflect slightly different developmental stages, with FBS/LIF conferring less naïve mouse ESCs than 2i/LIF conditions.

Other functional tests that can be used to assess the stringency and extent of naivety in different primate naïve PSCs, is whether the cells can tolerate complete ablation of epigenetic repressors like METTL3, DNMT1, DGCR8 and MBD3^4,66^ **(Fig. 3,4)**. Further, such tests might be useful for optimizing conditions that close the gap between mouse and human naive pluripotent cells isolated thus far **(Fig. 4, Box 1)**. Collectively, it will be informative to annotate different naïve and primed sates from any other species isolated thus far according to such criteria **(Fig. 3)**.

##### Box 1 A ‘dark side’ of naïve pluripotency?

With the development of naïve conditions and efforts to endow these cells with more features of naivety, one question that emerges is “how much naivety is needed and is there a dark side for permanently maintaining PSCs under naive conditions?”

Rodent ESCs expanded in 2i/LIF have an increased tendency for acquiring genomic abnormalities^41^, and it remains unclear whether these alterations occur as a by-product of nonspecific activity of small molecule inhibitors used^95,116^, or as a direct result of intrinsic molecular features of naïve pluripotency (e.g. increased endogenous retroviral elements (ERVs) activity, reduction in epigenetic repressive marks). One can envision a scenario where such features can be tolerated *in vivo* where this configuration exists for only 1-2 days, while prolonged *in vitro* expansion of this state increases the occurrence frequency of such unwanted damaging events.

The latter concern may also relate to safeguarding the integrity of DNA methylation and imprinting in naïve PSCs expanded *in vitro* over an extended period of time. Studies focusing on loss of DNA methylation upon transfer of mouse PSCs into 2i/LIF conditions have quantified methylation after 10-24 days of transfer, and have documented rapid global loss of DNA methylation accompanied with relative resistance of retrotransposons and imprinting regions to such demethylation^83^. However, it is unclear whether that latter described methylation state represents a final plateau that naive cells achieve, or whether carrying out 2i/LIF cultures further eventually leads to erosion in the relative resistance of such regions to demethylation. Indeed, methylation over imprinted genes and retrotransposons are partially, yet significantly, reduced in 2i/LIF conditions^83^.

If such effects are frequent, researchers will have to re-evaluate how to optimize applying naive conditions. One scenario might involve decreasing inhibitor levels as a mean to avoid excessive hypomethylation or other unwanted effects. An alternative scenario is to maintain cells in primed conditions and transfer them into naive conditions only for short time before initiation of differentiation.

### Implications and future directions

Yamanaka’s breakthrough of reprogramming somatic cells to pluripotency has provided the foundation for deeper sleuthing of pluripotent states and the understanding that pluripotent configurations can be rewired. The latter, feeds back and bares direct influence on issues associated with current hurdles and limitations related to human iPSC quality and characteristics **(Box 1,2)**.

##### Box 2 Potential implications on iPSC reprogramming.

Recent studies have implicated how certain epigenetic regulators in fact have opposing effects on naïve and primed murine PSC maintenance **(Fig. 4)**^66^. These findings might be relevant when comparing induction of pluripotency mechanisms in human vs. mouse, as human, but not mouse iPSCs, are typically reprogrammed in conventional/primed pluripotency conditions^117^. Consequently, some of the differences observed between human and mouse iPSC regulators might not be related to species differences but rather to the fact that distinct pluripotent states are being induced. Therefore, it would be imperative to expand human iPSC reprogramming regulator screens, and include different pluripotency conditions, as they might yield different outcomes.

Another implication of pluripotent state characteristics on reprogramming is whether naïve conditions might improve the quality of obtained iPSCs and facilitate loss of residual epigenetic memory and heterogeneity, when observed^118,119^. Similarly, subtle epigenetic differences in DNA methylation found between NT-ESCs and iPSCs generated from the same human donor cells^120^ might also be neutralized when deriving iPSCs in naïve conditions that mimic more closely epigenetic features of the ICM.

One of the most fascinating questions related to the naïve to primed pluripotency continuum is “why do these divergent pluripotent configurations actually exist? The latter is often accompanied by the pragmatic question of “Which cells are better to work with – naïve or primed?” In our opinion, as this phenomenon is deeply rooted in early embryonic development *in vivo,* it is likely that both configurations constitute essential and integral parts for safeguarding optimality and maximizing the benefits of multi-potency and lineage specification simultaneously. We hypothesize that naive pluripotency emerged as an epigenetic erasure state that renders pluripotent cells free from lineage and epigenetic restriction, while simultaneously making these cells relatively less responsive to signalling pathways that might interfere with establishing such a lineage neutral state. The induction of specification by morphogens may not be efficiently enforced during or immediately after establishing naïve pluripotency, without a short “delay period”. As such, the naïve pluripotency network is gradually resolved and becomes more receptive to inductive cues at the post-implantation stage, and PSCs get differentially patterned/primed according to their spatial localization, before overt somatic differentiation occurs.

At the functional level, it remains to be established whether using human naïve PSCs as a starting material, with or without a brief priming, would resolve hurdles currently faced in human *in vitro* PSC differentiation protocols: 1) Will human naïve PSCs yield increased consistency in differentiation among independent iPSC lines^110^? 2) Can naïve PSC conditions yield better quality cells in differentiation protocols when used as a starting material? 3) Can human naïve PSC facilitate the success of differentiation protocols that have not been conductive with conventional human PSCs? Encouraging support for the latter has recently been provided by showing the enhanced ability of human PSCs expanded in NHSM conditions (even in the absence of aPKCi) for undergoing *in vitro* differentiation into PGCs, a protocol that was inefficient from primed human PSCs^111^. The molecular rationale for evaluating the potential benefits highlighted above is that naïve pluripotency is more associated with cleansing of epigenetic repressive marks over regulatory regions, compared to primed cells^70,96^. This might enable more adequate activation of lineage specifiers during differentiation. Further, lineage biases in human primed PSCs are heavily associated with localized accumulation of repressive marks like DNA methylation^112^.

The recent advances in generating human naïve PSCs will continue to boost attempts to generate naïve-like PSCs from other species, and test same-species and inter-species embryo chimerism assays^113^. Cynomolgus monkey naïve ESCs derived in NHSM conditions gave rise to the first chimera competent non-human primate ESCs^114^. Developmentally advanced mouse embryos (E10.5-E17.5) with low chimerism levels were obtained following injection of naive human^70^ or monkey iPSCs^115^. These observations raise a variety of exciting challenges relating to defining what are the frequency, lineage preference and developmental quality of such integrated primate iPSC derived cells. Systematic efforts will be key to conclude whether humanized animal models^70^ might become relevant for disease modelling, studying human development or generate transplantable human organs^113^.

Continued breakthroughs in single cell technologies and applying them on different pluripotent cell types and embryonic samples will facilitate defining properties that are relevant for adequate functionality of PSCs. This will help set standards for optimal starting material for stem cell based therapeutics and research **(Box 1)**. It is expected that during this journey aiming at allowing scientists to better control cell fate by deconstructing this previously underappreciated complexity of pluripotency, proposed criteria and standards will likely be debated and revised.

<u>Display items</u> *(all will be subject to graphic editing by NRMCB after review)*

### Copyright permissions

- Figure 2,3,5a-b: Graphic figures are reused from our Nature Reviews Molecular Cell Biology Poster (http://www.nature.com/nrm/posters/pluripotency/index.html) as indicated in figure legends.
- Figure 5c is reused and modified from our Figure 3 in Manor et al. Curr. Opin. In Genetics & development

## Acknowledgements

J.H.H is supported by a generous gift from Ilana and Pascal Mantoux; the New Y ork Stem Cell Foundation (NYSCF), Flight Attendant Medical Research Institute (FAMRI), the Kimmel Innovator Research Award, the ERC-StG (StG-2011-281906) and ERC-PoC (PoC-2015-692945), Moross Cancer Institute, Israel Science Foundation – NFSC program, Morasha Biomed program, ICORE program, the ICRF Foundation, MINERVA fund, Helen and Martin Kimmel Institute for Stem Cell research, the Benoziyo Endowment fund, David and Fela Shapell Family Foundation INCPM Fund for Preclinical Studies, an HFSPO research grant. J.H.H. is a New York Stem Cell Foundation – Robertson Investigator. We thank W. Greenleaf and members of the Hanna lab for discussions. We apologize to those whose work could not be covered or directly cited due to space limitations.

## Further information

Online Web link sites:

Addgene plasmid repository: https://www.addgene.org/

ENCODE project: http://www.nature.com/encode/

Epigenome Roadmap project: http://www.nature.com/collections/vbqgtr

Mouse ES cell ChIP compendium: http://bioinformatics.cscr.cam.ac.uk/ES_Cell_ChIP-seq_compendium.html

Mouse ES Single cell RNA-seq resource – ESpresso: http://www.ebi.ac.uk/teichmann-srv/espresso/

CRISPR/CAS9 genome wide screen resource: http://genome-engineering.org/gecko/

Online poster by Nature Reviews Molecular Cell Biology on Pluripotent States: http://www.nature.com/nrm/posters/pluripotency/index.html

### Glossary

Primordial germ cells (PGCs): embryonic progenitor cells that give rise to germ cells in the gonads (sperm and oocytes).
Inner cell mass (ICM): the mass of cells inside the pre-implantation blastocyst that will subsequently give rise to the definitive structures of the fetus.
Embryonic stem cells (ESCs): *in vitro* expanded pluripotent cells that originate from the ICM.
Epiblast stem cells (EpiSCs): *in vitro* expanded pluripotent cells that originate from the postimplantation epiblast.
Embryonic germ cells: *in vitro* expanded pluripotent cells that are derived from embryonic PGCs.
Germ stem cells (GSCs): *in vitro* expanded pluripotent stem cells that originate from neonatal or adult testis derived spermatogonial stem cells.
Nuclear transfer: cloning of somatic cell derived nucleus and its introduction into a-nucleated host oocyte.
Induced pluripotent stem cells (iPSCs): *in vitro* generated pluripotent cells derived via ectopic expression of defined exogenous factors in somatic cells.
Naïve pluripotency: pluripotent state that resembles pre-implantation pluripotent configuration(s).
Primed pluripotency: pluripotent state that resembles to post-implantation embryonic configuration(s).
Ground state pluripotency: originally described as a state of pluripotency that is independent of exogenous activator signalling input or stimulation.
X inactivation: dosage compensation of X chromosome in female, where one of the X chromosomes gets epigenetically silenced.
Seed enhancers: subgroup of enhancers that are dormant in naive cells but become more active in primed pluripotent and somatic cells.
3i: Defined naïve pluripotency growth conditions combing 3 inhibitors (i) for MEK, FGF and GSK3 signalling.
2i/LIF: Defined naïve pluripotency growth conditions containing 2 inhibitors (i) for MEK and GSK3 together with LIF cytokine.
“Alternative 2i”: Defined naïve pluripotency growth conditions composed of 2 small molecule inhibitors for GSK3 and SRC pathways
LIF/MEKi/aPKCi: Defined naïve pluripotency growth conditions containing 2 inhibitors (i) for MEK and atypical PKC signalling, together with LIF cytokine.
FGF2/ACTIVIN A: Defined primed pluripotency growth conditions for mouse EpiSCs composed of recombinant FGF2 and ACTIVIN A cytokines.
GSK3i/IWR1: Defined primed pluripotency growth conditions for mouse EpiSCs composed of GSK3 pathway inhibitor and Tankyrase small molecule inhibitor, IWR1.
FGF2/IWR1: Defined primed pluripotency growth conditions for mouse EpiSCs composed of recombinant FGF2 and Tankyrase small molecule inhibitor, IWR1.

## Table of contents summary

Hanna and colleagues review recent advances on molecular underpinnings of alternative primed- and naïve-like pluripotent states isolated in rodents and in man. They highlight potential benefits and identify key unanswered challenges in this rapidly evolving fundamental topic.

## Online summary

1. Pluripotency is highly dynamic *in vivo* and evolves at different stages of pre- and postimplantation stages. However, the self-renewal aspect is a highly useful *in vitro* artificial phenotype endowed by culture conditions.
2. Different pluripotent cell <u>types</u> can be isolated *in vitro* from different sources and methods. The Pluripotent <u>state</u> assumed by the cultivated cells is dictated by the *in vitro* growth condition, rather than their cell of origin.
3. Naïve and primed pluripotent states can be functionally classified based on their ability or failure to maintain self-renewal of their pluripotent state upon MEK signaling inhibition, respectively.
4. Naïve and primed states of pluripotency can each represent a continuum of configurations rather than a fixed individual state. Within the naïve and primed pluripotent states, different degrees of naiveté or priming can be captured and annotated based on a variety of characteristics.
5. Human conventional pluripotent cells are primed, however they are not identical to mouse primed cells, and have certain naïve-like properties. *In vivo* differences likely underlie differences in growth requirements and characteristics of pluripotent cells isolated *in vitro* from mice and man.
6. The use of human naïve pluripotent growth conditions and cells might have great influence on iPSC/ESC quality and their differentiation competence, consistency and robustness.

## Author biographies

Leehee Weinberger M.Sc., has been a PhD student with Jacob H. Hanna at the Department of Molecular Genetics at the Weizmann Institute of Science, Rehovot, Israel for 4 years. She carried out her Masters work with Naama Barkai at the the same department. Her doctoral research focused on regulation and differentiation of human naïve pluripotent cells and germ cells.

Muneef Ayyash Ph.D., has been a staff researcher in Jacob H. Hanna lab at the Department of Molecular Genetics at the Weizmann Institute of Science Rehovot, Israel for 1 year. He carried his doctoral work with David Wallach at the same institute and did postdoctoral studies with Yinon Ben-Neriah at the Hebrew University, Jerusalem, Israel. His research currently focuses on WNT and HIPPO signaling modulation in promoting human naïve pluripotency.

Noa Novershtern Ph.D., has been a staff scientist and senior bioinformatician in Jacob H. Hanna lab at the Department of Molecular Genetics at the Weizmann Institute of Science, Rehovot, Israel for 5 years. She carried out her doctoral work with Nir Friedman at the Hebrew University, Jerusalem, Israel and Aviv Regev at the Broad Institute, Cambridge, USA. She focuses on dissecting epigenetic dynamics in pluripotency and reprogramming.

Jacob H. Hanna M.D. Ph.D., has been on the Faculty of the Department of Molecular Genetics at the Weizmann Institute of Science, Rehovot, Israel for 5 years, and is also a Robertson Investigator by the New York Stem Cell Foundation (NYSCF). He carried out his doctoral work with Ofer Mandelboim at the Hebrew University, Jerusalem, Israel and then did his postdoctoral studies with Rudolf Jaenisch at the Whitehead Institute for Biomedical Research - Massachusetts Institute of Technology, Cambridge, USA. His laboratory studies molecular mechanisms of stem cell reprogramming and differentiation. Hanna lab website: http://hannalabweb.weizmann.ac.il/

